# Atrophic astrocytes in aged marmosets present tau hyperphosphorylation, RNA oxidation, and DNA fragmentation

**DOI:** 10.1101/2023.02.09.527885

**Authors:** Juan D. Rodríguez-Callejas, Eberhard Fuchs, Claudia Perez-Cruz

**Affiliations:** Centro de Investigación y de Estudios Avanzados del Instituto Politécnico Nacional. Department of Pharmacology; German Primate Center, Leibniz-Institute of Primate Research, Göttingen, Germany.

**Keywords:** GFAP, 8-hydroxyguanosine, AT-100, non-human primate, entorhinal cortex, aging.

## Abstract

Astrocytes perform multiple essential functions in the brain showing morphological changes. Hypertrophic astrocytes are commonly observed in cognitively healthy aged animals, implying a functional defense mechanism without losing neuronal support. In neurodegenerative diseases, astrocytes show morphological alterations, such as decreased process length and reduced number of branch points, known as *astroglial atrophy*, with detrimental effects on neuronal cells. The common marmoset (*Callithrix jacchus*) is a non-human primate that, with age, develops several features that resemble neurodegeneration. In this study, we characterize the morphological alterations in astrocytes of adolescent (mean 1.75 y), adult (mean 5.33 y), old (mean 11.25 y), and aged (mean 16.83 y) male marmosets. We observed a significantly reduced arborization in astrocytes of aged marmosets compared to younger animals in the hippocampus and entorhinal cortex. These astrocytes also show oxidative damage to RNA and increased nuclear pTau (AT100). Astrocytes lacking S100A10 protein show a more severe atrophy and DNA fragmentation. Our results demonstrate the presence of atrophic astrocytes in the brains of aged marmosets.

## 1. Introduction

The common marmoset (*Callithrix jacchus*) is a small New-World primate with high genetic homology to humans and a shorter life span than Old-World primates (Abbott and Barnett, 2003). Marmosets are widely used in biomedical research due to their small size (20-30 cm and 250-600 g) and high reproductive capacity (Okano et al., 2012). For aging research, marmosets are positioned as ideal non-human primate (NHP) model as they show signs of aging around 8 years of age, while their life span in captivity reaches up to 20 years (Kramer and Burns, 2019). Several groups have documented cognitive impairment in aged marmosets: working memory impairment is observed in old marmosets compared to younger subjects (De Castro and Girard, 2021; Rothwell et al., 2022; Workman et al., 2018), and delay response strategy is detectable in adult marmosets (Sadoun et al., 2019, 2015). Also, old marmosets develop amyloid plaques in the cortex and tau hyperphosphorylation (pTau) in cortical and hippocampal regions. Notably, pTau can be detected in adolescent marmosets, significantly increasing with aging (Rodriguez- Callejas et al., 2016). Dystrophic microglia, iron accumulation, and increased oxidative stress are features of old and aged marmosets (Rodríguez-Callejas et al., 2019). Thus, the common marmoset has been positioned as an ideal model for understand the etiological factors associated with aging and neurodegeneration (t’Hart et al., 2012; Rodriguez- Callejas et al., 2016; Ross and Salmon, 2018; Rodríguez-Callejas et al., 2019; Sadoun et al., 2019).

Astrocytes are star-shaped glial cells with radial processes that play essential functions as blood-brain barrier (BBB) formation and maintenance (Abbott et al., 2006; Daneman and Prat, 2015; Janzer and Raff, 1987), ionic environment regulation (Anderson and Swanson, 2000; Sattler and Rothstein, 2006; Seifert et al., 2006; Simard and Nedergaard, 2004; Strohschein et al., 2011), control of neurogenesis and glycogen storage (Brown et al., 2005; Brown and Ransom, 2007; Matsui et al., 2017), neurometabolic uncoupling (Magistretti, 2006), iron-induce antioxidant protection (Hoepken et al., 2004; Oide et al., 2006; Regan et al., 2002), among others. When the brain tissue is damaged, astrocytes exhibit hypertrophy (Hol and Pekny, 2015; Kimelberg, 2004), and alter gene expression of glial fibrillary acidic protein (GFAP) resulting in a *reactive state* (Sofroniew, 2009). Previous reports indicates that astrocytes of aged rats showed an enhanced expression of GFAP and hypertrophic phenotype, similar to data from old humans, and senescence-accelerated animal models (Clarke et al., 2018; Cotrina and Nedergaard, 2002; Hol and Pekny, 2015; Kohama et al., 1995; Nichols et al., 1993; Rodríguez et al., 2014; Rozovsky et al., 1998; Woodruff-Pak, 2008; Wu et al., 2005; Yoshida et al., 1996). In aged mice, this reactive phenotype is characterized by an increase processes surface, volume, and somata. In contrast, in the entorhinal cortex (ENT) of aged mice, astrocytes present smaller processes, a condition named *atrophic astrocytes* (Rodríguez et al., 2014). In cognitive-healthy aged humans and NHPs, atrophic astrocytes can be detected in *substantia nigra pars compacta* and the midbrain, respectively (Jyothi et al., 2015; Kanaan et al., 2010). However, in mouse models of AD, astroglial atrophy is heavily observed not only in ENT, but also in DG, CA1, and medial prefrontal cortex (Beauquis et al., 2013; Kulijewicz-Nawrot et al., 2012; Olabarria et al., 2010; Verkhratsky et al., 2015). Therefore, atrophic astrocytes in vulnerable brain regions may be related to a neurodegenerative process. Marmosets develop several features associated with neurodegeneration during aging (i.e., amyloid plaques, pTau, dystrophic microglia, oxidative damage) (Geula et al., 2002; Maclean et al., 2000; Palazzi et al., 2006; Philippens et al., 2016; Ridley et al., 2006; Rodriguez-Callejas et al., 2016; Rodríguez-Callejas et al., 2019; Sharma et al., 2019), but up to day there is no information about the presence of atrophic astrocytes in this NHP. Therefore, we aimed to determine the phenotypic alterations in astrocytes of the hippocampus and ENT in the marmoset during aging.

For this, we evaluated morphological alterations in GFAP+ astrocytes by the Sholl analysis in ENT and hippocampal region (CA1, CA2-CA3) of male adolescence, adult, old, and aged marmosets. We also determined the number of astrocytes with oxidative damage to RNA, pTau, and S100A10 protein (S100A10+) by double-immunolabeling. We found an enhanced number of reactive astrocytes (defined by an increased process length, volume, and branch points) in adult and old marmosets compared to adolescents, featuring a reactive phenotype. However, in aged marmosets, these features were significantly decreased in all regions analyzed, showing an atrophic phenotype. In addition, old and aged marmosets showed an enhanced number of astrocytes with RNA oxidation and nuclear pTau. Furthermore, in old subjects, astrocytes with shorter processes length and branching points were S100A10-, a marker of neuroprotective astroglial phenotype. S100A10-astrocytes presented fragmentation of DNA (TUNEL staining) and were clearly atrophic. Thus, our results show that marmosets develop astroglia atrophy in the hippocampus and ENT during the aging process. This data offers further evidence that marmosets develop several features of a neurodegenerative process with aging.

## 2. Methods

### 2.1 Subjects

Laboratory-bred common marmosets (*Callithrix jacchus*) were housed at the German Primate Center, Göttingen, Germany, under standard conditions complying with the European Union guidelines for the accommodation and care of animals used for experimental and other scientific purposes (2007/526/EC). All animal experiments were performed in accordance with the German Animal Welfare Act, which strictly adheres to the European Union guidelines (EU directive 2010/63/EU) on the use of NHP for biomedical research. Experienced veterinarians and caretakers constantly monitored the animals. Animals did not present neurological disorders or other injuries that can cause trauma to the central nervous system.

### 2.2 Tissue preparation

Brains of male marmosets of different ages were used in the current study: Four adolescents (A: mean age 1.75 ± 0.18 years old), three adults (Ad: mean age 5.33 ± 0.88 years old), five old (O: mean age 11.25 ± 0.70 years old), and three aged (Ag: mean age 16.83 ± 2.59 years old) individuals based on previous age classification (Abbott and Barnett, 2003). Animals were anesthetized with an i.p. injection (0.1 ml/100 g body weight) of GM II (ketamine, 50 mg/ml; xylazine 10mg/ml; atropin 0.1 mg/ml) and after loss of consciousness they received an i.p. injection of ketamine (400 mg/kg body weight). Bodies were transcardially perfused with cold (4 °C) saline (0.9 % NaCl) for 5 min. Subsequently, for fixation of the brains, cold (4 °C) 4 % paraformaldehyde (PFA) in 0.1 M phosphate buffer, pH 7.2, was infused for 15 min. The brains were removed and post-fixed in fresh 4 % PFA at 4 °C, where brains were stored until sectioning. Before sectioning, tissue was washed thoroughly with 0.1 M phosphate-buffered saline (PBS:

0.15 M NaCl, 2.97 mM Na_2_HPO_4_-7H_2_O, 1.06 mM KH_2_PO_4_; pH 7.4), and immersed in 30 % sucrose in PBS at 4°C four days before sectioning. Coronal sections (40 μm thick) were obtained using a sliding microtome (Leica RM2235) and we prepared series every 6th section (at intervals of 240 μm) from the hippocampal formation (Bregma +8.00 mm to +0.80 mm) according to Paxinos et al.(2012). Sections were immediately immersed in cryoprotectant solutions for immunofluorescence [300 g sucrose; 10 g polyvinyl- pyrrolidone (PVP-40); 500 mL of 0.1M PBS and 300 mL ethylene glycol, for 1 L] and stored at -20°C until use.

### 2.3 Double labeling immunofluorescence

For double labeling of astrocytes with AT100 and GFAP, brain sections were pretreated with formic acid for 15 min and with citrate buffer 20X at 94 °C for 10 min. To reduce autofluorescence and background due to PFA fixation, sections were incubated with 1% sodium borohydride (NaBH_4_) in PBS-1X for 10 min. Then, sections were rinsed with 0.5 % PBS-Tween20 twice for 3 min. To block potential nonspecific antibody binding, sections were incubated for 30 min using a solution containing 2 % donkey serum, 50 mM glycine, 0.05 % Tween20, 0.1 % TritonX-100, and 0.1 % BSA diluted in PBS. Primary antibody anti-GFAP (1:300, goat/IgG, Abcam, Ab53554) was incubated in an antibody signal enhancer (ASE) solution (Flores-Maldonado et al., 2020; Rosas-Arellano et al., 2016), consisting of 10 mM glycine, 0.05 % Tween20, 0.1 % TritonX-100 and 0.1 % hydrogen peroxide in PBS, and left overnight at 4 °C. For double labeling, GFAP antibody was incubated with 8-hydroxyguanosine (8OHG, marker of RNA oxidation) (1:10000, mouse/IgG, Abcam, ab62623), AT100 (1:500, mouse/IgG, Thermo Scientific, MN1060), and S100A10 (1:250, mouse/IgG, Invitrogen, PIMA515326) primary antibodies. On the next day, sections were washed with 0.5 % PBS-Tween20, and thereafter, incubated with secondary antibodies ALEXA647 anti-goat (1:500, donkey/IgG, Jackson ImmunoResearch, 705-605-147) and ALEXA488 anti-mouse (1:500, donkey/IgG, Jackson ImmunoResearch, 715-545-150), diluted in 0.1 % PBS- Tween20 for 2 hours at RT. All sections were incubated with DAPI (1:1000, Affymetrix) in 0.2 % PBS-triton for 30 min. To reduce lipofuscin autofluorescence, brain sections were incubated in 0.1% Sudan black (Sigma) for 15 min. Finally, sections were washed and mounted on glass slides with mounting medium VectaShield (Vector Laboratories).

### 2.4 Double labeling immunofluorescence and TUNEL protocol

To detect DNA fragmentation in GFAP+ astrocytes an *in situ* cell death detection kit (Roche, 11684795910) was used. Before TUNEL reaction, brain sections were incubated with 1% sodium borohydride (NaBH_4_) in PBS-1X for 10 min. Then, sections were permeabilized with 0.3% PBS-triton for 20 min. For TUNEL reaction, sections were incubated in the mixture of label solution and enzyme solution at 37 °C for 1 h. Sections were rinsed with PBS-1X for 10 min. Thereafter, double labeling immunofluorescence for GFAP and S100A10 was performed as described in 2.3.

### 2.5 Image acquisition

Images were obtained by a confocal microscope (Leica TCS-SP8) equipped with Diode (405 nm), OPSL (488 nm), OPSL (552 nm), and Diode (638 nm) laser. Brain regions were imaged performing optical scanning with 732 gain, -2.8 offset, and 1.0 UA pinhole diameter. For double labeling (GFAP versus 8OHG, AT100, or S100A10) and Sholl analysis images were acquired with a 63X objective. All confocal images were obtained as z-stacks of single optical sections. Stacks of optical sections were superimposed as a single 2D image by using the Leica LASX software. We captured images from different regions of the hippocampus (DG, CA3, and CA2-CA1) and ENT according to the marmoset brain atlas (Paxinos et al., 2012).

### 2.6 Sholl analysis

To quantify the length, volume, and number of branching points of astrocytic processes, a Sholl analysis was performed by using NeuronStudio software (Canchi et al., 2017). For this analysis, we captured 3 images per brain region analyzed (DG, CA3, CA2-CA1, and ENT), per subject.

In every captured image, ten astrocytes were analyzed, given a total of 1440 astrocytes analyzed. All the images were captured with the following parameters: optical gain 732, offset – 3.0, and 1.0 AU pinhole diameter. For Sholl analysis, image z-stacks were imported to NeuronStudio software for the reconstruction of astrocytes processes. Sholl analysis is based on the generation of concentric spheres radiating from the cell center and at each sphere, the length, volume, and the number of branch points are quantified.

In our experiment, we run a Sholl analysis with every concentric sphere 1 µm larger than the previous one. To calculate the total length, total volume, and the number of branch points, we sum the values of the respective parameter (length, volume, or branch points) obtained in all the concentric spheres (radius) of the astrocyte analyzed.

### 2.7 Sholl analysis of S100A10 positive and negative astrocytes of aged marmosets

Microglia can be classified into two major subtypes (proinflammatory and phagocytic) (Cameron and Landreth, 2010; Franco and Fernández-Suárez, 2015; Kabba et al., 2017; Orihuela et al., 2016; Tang and Le, 2016). Previous data in marmosets have shown that aging is accompanied by an increased number of phagocytic-ameboid dystrophic microglia (Rodríguez-Callejas et al., 2019). Dysfunctional phagocytic microglia renders neuronal cells vulnerable to further damage. As we observed major astrocyte atrophy in aged marmosets, we decided to determine whether those cells are S100A10+, a specific maker of neuroprotective phenotype (Clarke et al., 2018). We captured three images per brain region (DG, CA3, CA2-CA1, and ENT) on each aged subject. Two S100A10+ and two S100A10- astrocytes were analyzed per image, given a total of six astrocytes of each type per region, on each subject. Sholl analysis and quantifications of length, volume, and branch points were performed according to 2.6.

### 2.8 Quantification of GFAP fluorescence intensity

The quantification of GFAP fluorescence intensity was performed in 1440 astrocytes. This analysis was performed according to previous studies (Flores-Maldonado et al., 2020; Rosas-Arellano et al., 2016). Using the ImageJ “free hand” function the cytoplasm and processes of the astrocytes were outlined. Then, we quantified the fluorescence intensity of the selected area. The background signal was subtracted from the positive signal to obtain the relative intensity of the GFAP signal.

### 2.9 Number and percentage of 8OHG+/AT100+ astrocytes per brain region

To determine the number of astrocytes with damage to the RNA and tau hyperphosphorylation, we double-labeled GFAP+ astrocytes with 8OHG and AT100. Three images were captured on each region analyzed (DG, CA3, CA2-CA1, and ENT) with a 63x objective. All the images were captured with the following parameters: GFAP (optical gain 732 and offset – 3.0), 8OHG (optical gain 680, offset – 1.0), and AT100 (optical gain 680, offset – 3.0). The quantification of GFAP+/8OHG+/AT100+astrocytes was performed using ImageJ software (Plugins---Analyze---Cell counter). The sum of double-labelled astrocytes in a certain region was divided by the total area analyzed. The total area analyzed was calculated by multiplying the area of a 63x image (0.034 mm^2^) by three (the number of images captured on every hippocampal region), resulting 0.10 mm^2^ in all regions analyzed.

To calculate the percentage of 8OHG+/AT100+/GFAP+ astrocytes, the sum of 8OHG+/AT100+/GFAP+ astrocytes was multiplied by 100, and the final product was divided by the total amount of astrocytes in the region analyzed (the sum of 8OHG+/AT100+/GFAP+ astrocytes plus the astrocytes with no labeling for 8OHG or AT100, respectively).

### 2.10 Statistical analysis

Statistical analysis was performed by one-way ANOVA, followed by a Tukeýs as posthoc test, except for the quantification of astrocytic processes length using Sholl analysis where a multiple-t test followed by Holm-Sidak analysis by use of GraphPad Prism 6.0 software. Differences were considered statistically significant when p ≤ 0.05. Data are presented as means ± S.E.M.

## 3. Results

### 3.1 The number of GFAP+ astrocytes did not decrease with age in the common marmoset

We quantified the number of GFAP+ astrocytes in DG, CA3, CA2-CA1, and ENT of adolescent, adult, old, and aged marmosets. There were no significant differences in the number of GFAP+ astrocytes in any brain region analyzed, except for a significant decrease in adult subjects compared to adolescents in DG and in ENT of aged subjects compared to old marmosets (figure 1). This result indicates that the population of GFAP+ astrocytes remains similar during aging in the hippocampus and ENT of the marmoset. However, in adult and old animals, GFAP+ astrocytes had longer processes, and their labeling intensity was greater than in adolescent and adult individuals. Contrary, astrocytes of aged marmosets showed smaller process length and lower label intensity than astrocytes from the other age groups (figure 2). These morphological alterations suggest reactive astrogliosis in adult and old marmosets but astroglial atrophy in aged marmosets. To better characterize these morphological alterations in astrocytes, we used Sholl analysis.

**Figure 1.**
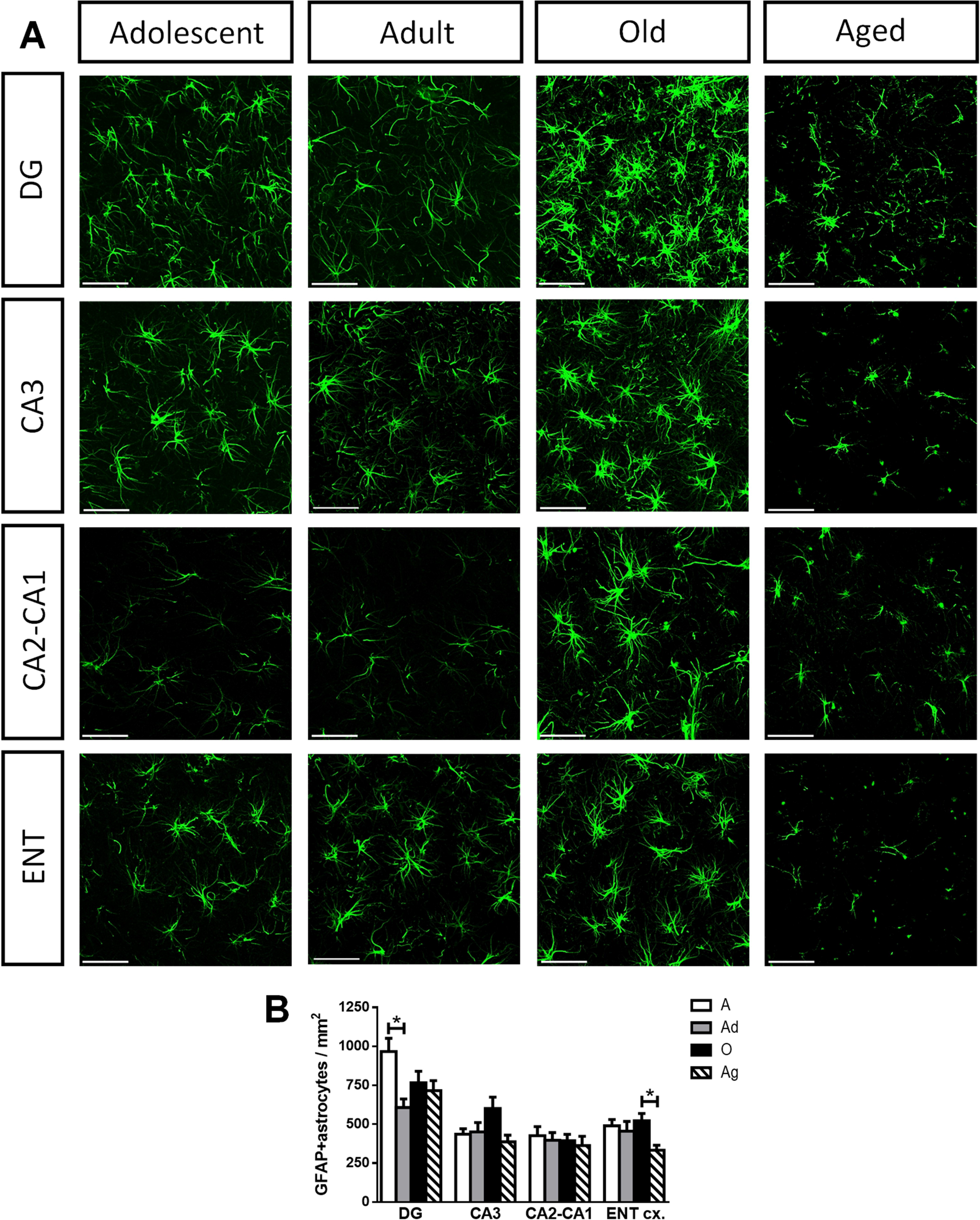
GFAP+ astrocytes the brain of marmosets of different ages in entorhinal cortex and hippocampus (DG, CA1, CA2-CA3). A. Representative photomicrographs of GFAP+ astrocytes in adolescence, adult, old and aged marmosets in selected brain regions. Scale bars 40 µm. **B. Quantification of GFAP+ astrocytes per brain region**. The number of GFAP+ astrocytes did not change with age in CA3, CA2-CA1 regions. In DG, there was a significant decreased in the number of GFAP+ in adults compared to adolescents. In ENT, aged marmoset had a decreased number of GFAP+ astrocytes than old subjects. Data represent means ± S.E.M. One-way ANOVA, Tukey post hoc analysis. *p < 0.05.

**Figure 2.**
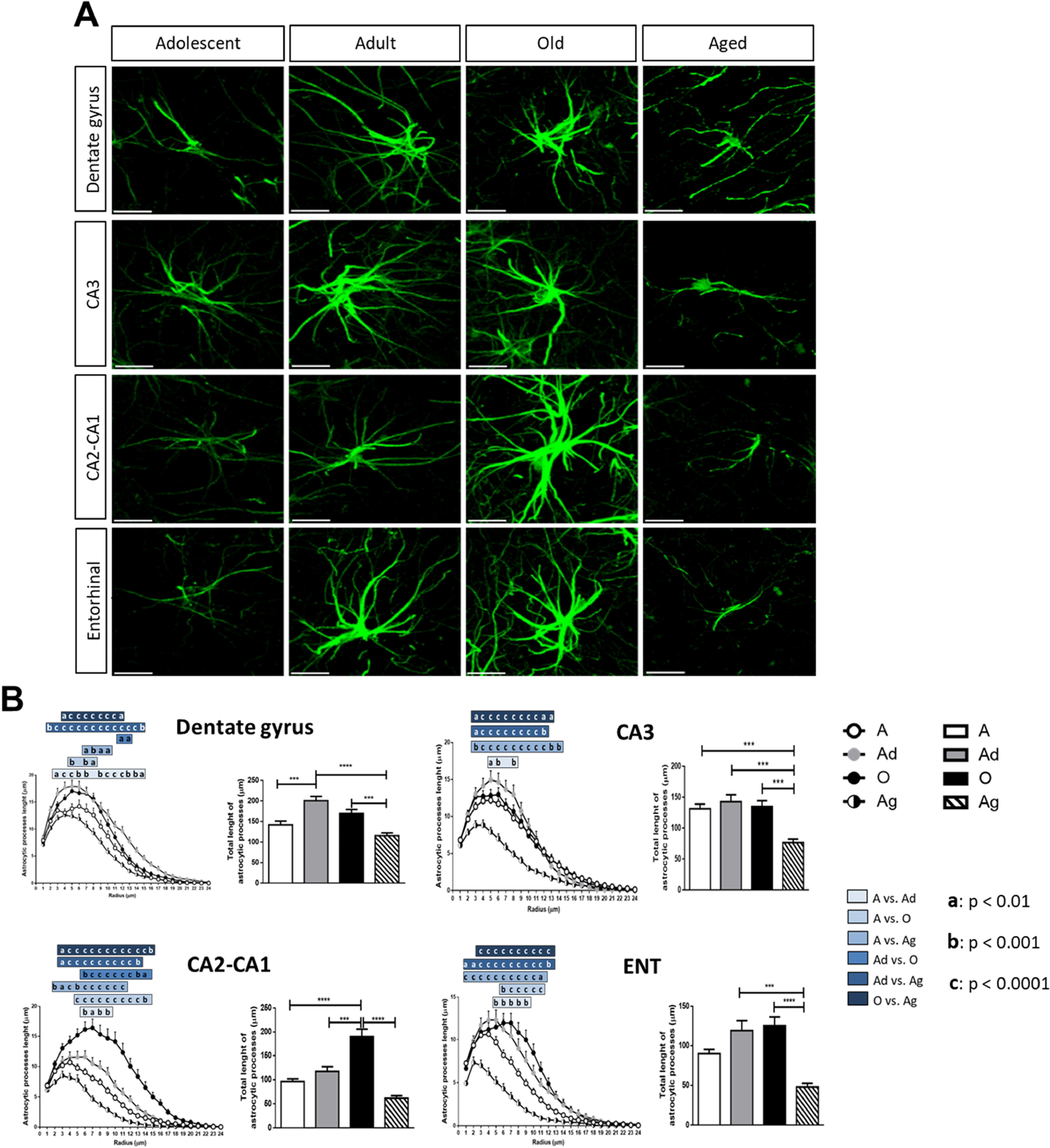
Morphological changes in astrocytes during the aging process in marmosets. A) Representative photomicrographs of GFAP+ astrocytic of adolescent, adult, old and aged marmosets in the brain regions analyzed. Note that in dentate gyrus and ENT, astrocytes of adult and old marmosets showed longer astrocytic processes than astrocytes of adolescent and aged subjects. In CA3, astrocytes of adolescent, adult and old marmosets showed longer astrocytic processes than astrocytes of aged subjects. In CA2-CA3, astrocytes of old marmosets showed longer astrocytic processes than astrocytes of all other age groups. Scale bars 10 µm. **B)** Sholl analysis of GFAP+ astrocytes. Astrocytic process length (APL) and total astrocytic process length (total APL) in the hippocampus and ENT of marmosets at different ages. DG showed a significantly increased APL in adults (radius 3-15, except 8) and old (radius 5, 7 and 8) marmosets with respect to adolescent. In aged marmosets, APL decreased with respect to any other age group (adolescents, radius 7-10; adults, radius 2-15; old, radius 4-12). Total APL was increased in adult marmoset compared to adolescents. In aged marmosets, total APL showed a significant reduction with respect to adult and old marmosets. CA3 region showed a significantly decreased APL in aged marmoset compared to any other age groups (adolescents: radius 3-14; adults, radius 3-12; old, radius 3-13). Aged marmosets showed a significant decreased of total APL than the rest of the groups (all p < 0.001). CA2-CA1. APL increased in old marmosets compared to adolescents (radius 5-14), adults (radius 6-14) and aged (radius 3-15) marmosets. In aged marmosets, APL decreased compared to any other age groups (adolescents, radius 2-11; adults, radius 3-13; old, radius 3-15). Old subjects present a greater total APL compared to adolescents, adults and aged marmosets. ENT. APL significantly increased in adults (radius 5-9) and old (radius 6-11) marmosets with respect to adolescents. In aged subjects, APL decreased compared to any other age groups (adolescents, radius 1-12; adults, radius 1-13; old, radius 3-12). Aged subjects showed a major decreased in total APL compared to adults and old marmosets. Data represent means ± S.E.M. For radial analysis of APL, a multiple t-test followed by Holm-Sidak post hoc analysis was performed with **p < 0.01, ***p < 0.001; ****p < 0.0001. For quantifications of total APL, one-way ANOVA followed by Tukey post hoc analysis was performed. a (**p < 0.01), b (***p < 0.001) and c (****p < 0.0001)

### 3.2 Astrocytic length, volume, and branching points decreased in different brain regions in aged marmoset

Using Sholl analysis we quantified the length, volume, and branching points of astrocytic processes of adolescent, adult, old, and aged marmosets, in DG, CA3, CA2-CA1, and ENT. We quantified the astrocytic processes length (APL) in adolescents (white circles curves), adults (grey circles curves), and old marmosets (black circles curves) (figure 2B and Supl. table 1). APL reaches a maximum between radius 4-7. This starts to decrease around radius 6-8 until reaching zero in radius 24. However, in aged marmosets (white and black curves) the maximum amount of APL is reached between radius 2-4, and then it rapidly decreased (figure 2B).

In DG, astrocytes of adults and old marmosets present a significant increase of APL compared to adolescents (adolescents vs. adults: from radius 3 to 7 and from 9 to 15; adolescents vs. old: radius 5, 7 and 8. Suppl. table 1); however, in astrocytes of aged marmoset APL was significantly decreased compared to all other age groups (aged vs. adolescents: from radius 7 to 10; aged vs. adults: from radius 2 to 15; aged vs. old: from radius 4 to 12. Suppl. table 1). In CA3, adolescents, adults, and old marmosets present similar APL values (significant differences in radius 5, 6, and 8 of adolescents vs. adults. Suppl. table 1), while astrocytes of aged marmosets showed a significant decreased of APL compared to all other age groups (aged vs. adolescents: from radius 3 to 14; aged vs. adults: from radius 3 to 12; aged vs. old: from radius 3 to 13. Suppl. table 1). In CA2- CA1, astrocytes of adolescents and adults have similar amounts of APL (significant differences from radius 6 to 9. Suppl. table 1). However, astrocytes of old marmosets showed a higher APL compared to adolescents and adults (old vs. adolescents: from radius 5 to 14; old vs. adults: from radius 6 to 14. Suppl. table 1). In aged marmosets, the APL significantly decreased compared to adolescents, adults, and old (aged vs. adolescents: from radius 2 to 11; aged vs. adults: from radius 3 to 13; aged vs. old: from radius 3 to 15. Suppl. table 1). In ENT, adult and old marmosets had a significantly increased of APL compared to adolescents (adolescents vs. adults: from radius 5 to 9; adolescents vs. old: from radius 6 to 11. Suppl. table 1). Astrocytes of aged marmoset present a significantly decreased APL compared to all other age groups (aged vs. adolescents: from radius 1 to 12; aged vs. adults: from radius 1 to 13; aged vs. old: from radius 3 to 12. Table 1).

The total length of the astrocytic process (total APL), that is the sum of APL from the twenty-four radius for each astrocyte, was also quantified. In DG and ENT, the total APL increased in adults and old subjects compared to adolescents and it significantly decreased in aged subjects. In CA3, adolescents, adults, and old marmosets have similar total APL, but in aged subjects, it was significantly decreased compared to all other age groups. In CA2-CA1, old marmosets present a significantly higher total APL compared to all other age groups (figure 2B).

Sholl analysis provides information as the caliber of the processes (volume) and the number of branch points. We analyzed the astrocytic processes volume (APV) in all regions studied. APV showed a similar trend to APL (figure 3C). In DG and ENT, the APV of adults and old subjects significantly increased with respect to adolescents and significantly decreased in aged subjects. In CA3, adolescent, adult, and old marmosets present similar APV. However, in aged marmosets, APV significantly decreased compared to all other age groups. In CA2-CA1, old marmosets have significantly higher APV compared to all other age groups.

**Figure 3.**
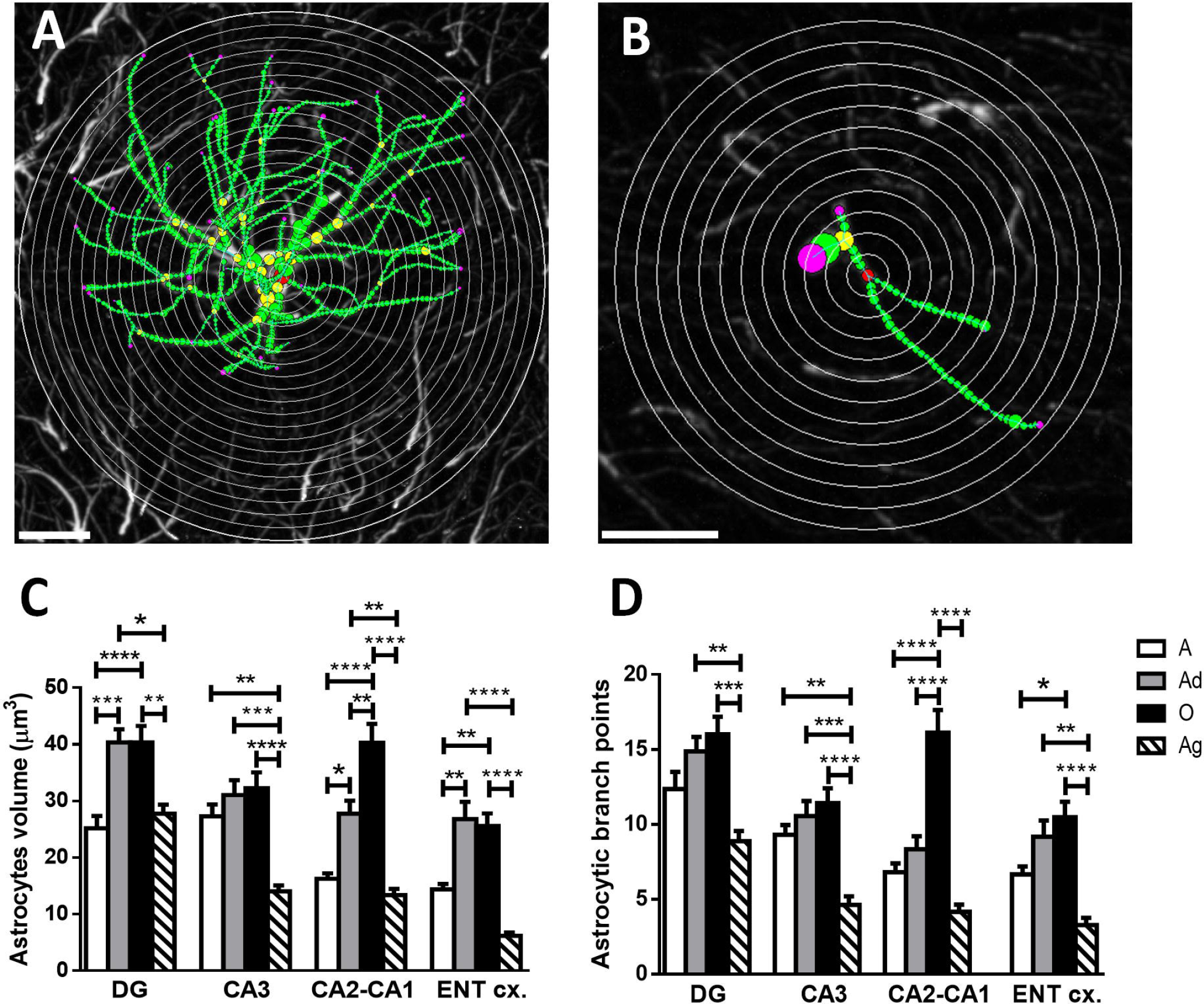
Volume and branching points of astrocytic processes in different brain regions of marmoset during aging. Representative images of the Sholl analysis by concentric spheres of an hypertrophic astrocyte (**A**) and an atrophic astrocyte (**B**). Processes’ ends are represented as pink circles and branch points as yellow circles. Scale bars 10 µm. **C)** Quantification of astrocytic process volume (APV). The volume of astrocytic processes of adult and old marmosets increased compared to adolescent subjects in DG, CA2-CA1 and ENT. However, in aged marmosets, APV was significantly reduced in all regions compared to adult and old marmosets. In CA3, astrocytes of aged marmosets showed a significantly decreased APV compared to adolescents, adults and old subjects. **D)** Quantification of astrocytic branching points (ABP). In the four regions analyzed the number of ABP showed a tendency to increase with age. However, it was only significant in CA2-CA1 where old marmoset had an increased ABP compared to adult and adolescent animals. In aged marmosets, ABP decreased in all regions compared to the other age groups. Data represent means ± S.E.M. One-way ANOVA, Tukey post hoc analysis. *p < 0.05; **p < 0.01; ***p < 0.001; ****p < 0.0001.

The astrocytic branch points (ABP, figure 3D) in DG, CA3, CA2-CA1, and ENT, were also analyzed. ABP tended to increase from adolescents to old marmosets. Nonetheless, ABP of aged subjects was significantly decreased with respect to all the other groups. CA2-CA1 showed a different trend since in this region old subjects had a significantly higher ABP compared to all other groups. These results indicate that despite the number of GFAP+ astrocytes does not increase during aging, astrocytes develop alterations in the length of their processes, volume, and branching complexity.

Adults and old marmosets present an increased length and volume of the astrocytic processes compared to adolescents, probably reflecting a reactive role against age-related insults. However, in aged marmosets, the length/volume of the processes was decreased (as an indicator of astrocytic atrophy). The loss of branching complexity may impact the functions of these cells.

### 3.3 Decreased GFAP-fluorescence intensity in astrocytes of aged marmosets

Besides of morphological alterations in astrocytes of aged marmosets, we also observed alterations in the fluorescence intensity (suppl. figure 1A). In DG and ENT, adolescent, adult, and old marmosets have similar levels of GFAP-fluorescence intensity (GFAP-FI) (suppl. figure 1B). Contrary, aged marmosets showed a significantly decreased GFAP-FI concerning adolescents and old subjects. In CA3 and CA2-CA1, GFAP-FI significantly increased in old subjects with respect to adolescents and adults (supplementary figure 1B); However, in aged marmosets, GFAP-FI significantly decreased compared to old subjects. The decrease of the GFAP-FI in astrocytes of aged marmosets may suggest a decreased expression of GFAP protein, the main component of the astrocyte cytoskeleton. However, as all the tissue available for this study is on PFA, we could not determine GFAP expression of its protein levels.

### 3.4 RNA oxidation in astrocytes increased in the hippocampus of aged marmoset

In a previous study, we observed an increase in RNA oxidation (8OHG) in aged marmoset brain (Rodríguez-Callejas et al., 2019). Neurons labeled with 8OHG+ were lightly detected in adolescent subjects but were significantly increased with aging. However, 8OHG+ microglia were only observed in aged subjects. In this study, we noticed that 8OHG+ astrocytes were almost absent in adolescent and adult marmosets. Nonetheless, old and aged subjects present abundant 8OHG+ astrocytes (figure 4). Quantification of 8OHG+ astrocytes per area shows an increase of 8OHG+ astrocytes in old and aged marmosets compared to adolescents and adults (figure 4B). This dramatically increase of RNA oxidation in astrocytes of aged marmosets was confirmed when we calculated the percentage of 8OHG+ astrocytes (figure 4C). In aged subjects, the percentage of 8OHG+ astrocytes reached approximately 50 to 80 % of the total astrocytes labeled with GFAP depending of the brain region (DG: 49.18 %; CA3: 67.59%; CA2-CA1: 80.89%; ENT: 71.46%), whereas in adolescent (DG: 2.96 %; CA3: 3.37 %; CA2-CA1: 8.58 %; ENT: 9.38 %) and adults (DG: 4.57 %; CA3: 4.49 %; CA2-CA1: 3.92 %; ENT: 2.08 %) this percentage was lower.

**Figure 4.**
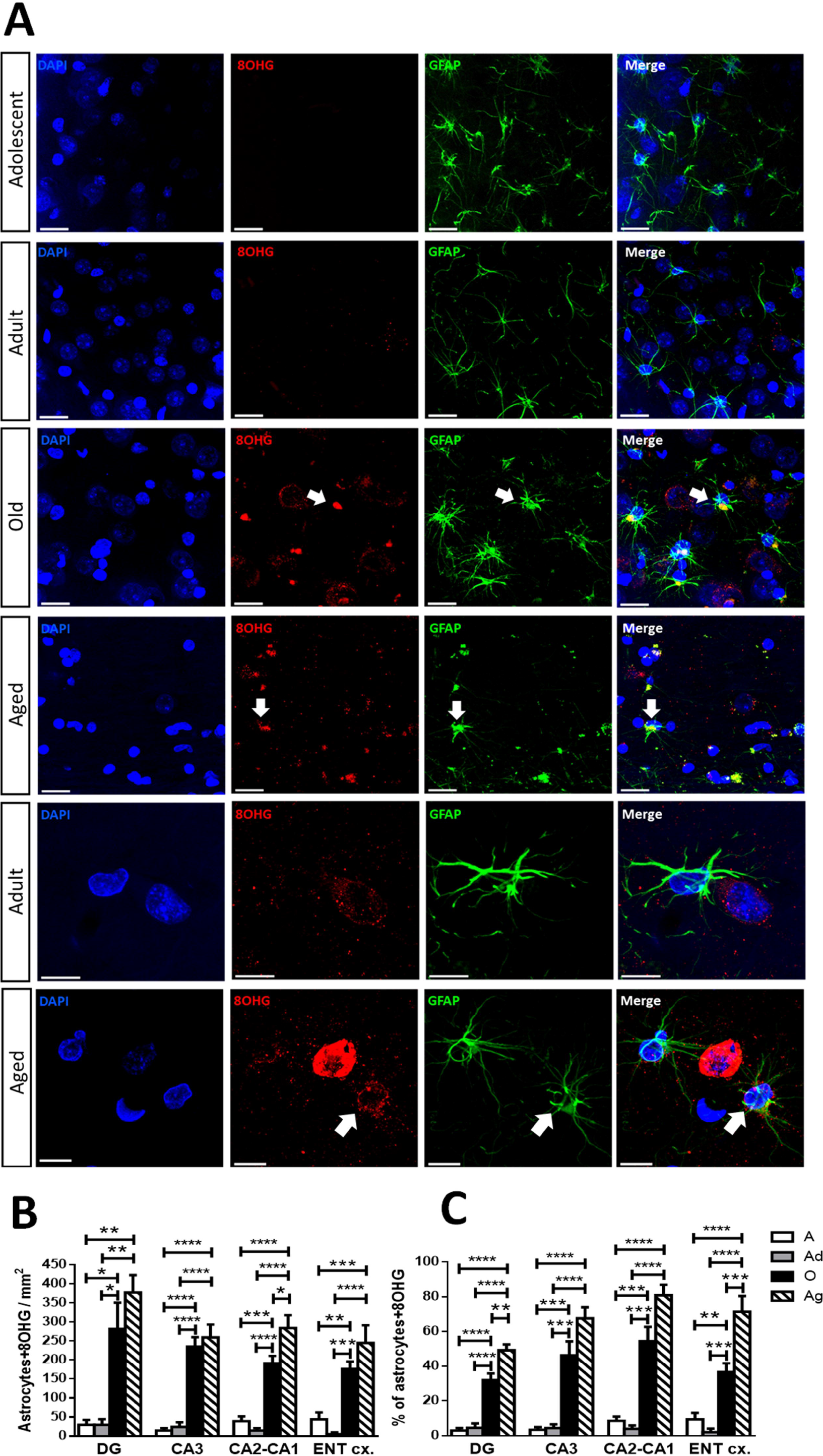
RNA oxidation in astrocytes of marmosets. A) Representative photomicrographs of astrocytes (GFAP, green) double labeled with 8OH (red, RNA oxidation marker 8OHG in brain sections of adolescent, adults, old and aged marmosets in CA3. Scale bar 20 µm. Last two panels show a higher magnification of astrocytes in adult and aged marmosets. In the aged subject, two 8OHG+ astrocytes are flanking an 8OHG+ cell, but one astrocyte also show 8OHG-labeling in the cytoplasm (white arrow). DAPI was used as a nuclear counterstain. Scale bar 10 µm. **B)** Number of astrocytes double labelled with 8OHG+ (RNA oxidation) in different brain regions. The number of 8OHG+ astrocytes are significantly increased in old and aged marmosets compared to adolescents and adults, in all regions analyzed. **C)** Percentage of astrocytes labeled with 8OHG. Old and aged marmosets show a higher percentage of 8OHG+ astrocytes compared to adolescent and adult subjects. CA2-CA1 was the region with the highest increase of 8OHG+ astrocytes in old and aged marmosets, followed by CA3 and ENT. DG was the region with the lowest percentage of 8OHG+ astrocytes. Data represent means ± S.E.M. One-way ANOVA, Tukey post hoc analysis. *p < 0.05; **p < 0.01; ***p < 0.001; ****p < 0.0001.

### 3.5 Nuclear hyperphosphorylated tau increased in astrocytes of marmoset with aging

In a previous study, we detected pTau in the residues Thr212 and Ser214 using AT100 antibody. AT100 label were located in the nuclear compartment of cells (Rodriguez-Callejas et al., 2016). To determine if the morphological alterations observed in atrophic astrocytes were related to the presence of pTau, we performed a double labeling GFAP / AT100 in all brain regions analyzed. As shown in figure 5, adolescent marmosets had few astrocytes with AT100. In adult, old, and aged marmosets, this number increased in the four regions analyzed. The quantification of AT100+astrocytes per area shows an increase of AT100+ astrocytes in DG, CA3, and CA2-CA1 of adult and old marmosets compared to adolescents (not significant). However, in aged marmosets AT100+ astrocytes decreased (not significant) compared to adult and old subjects. In ENT, AT100+ astrocytes significantly decreased in old and aged compared to adult subjects (figure 5B). We also calculated the percentage of AT100+ astrocytes per brain region analyzed (figure 5C). In the four regions analyzed, the percentage of AT100+ astrocytes increased in adult, old, and aged animals compared to adolescents. These results demonstrate that the presence of pTau increase with age in astrocytes of marmosets.

**Figure 5.**
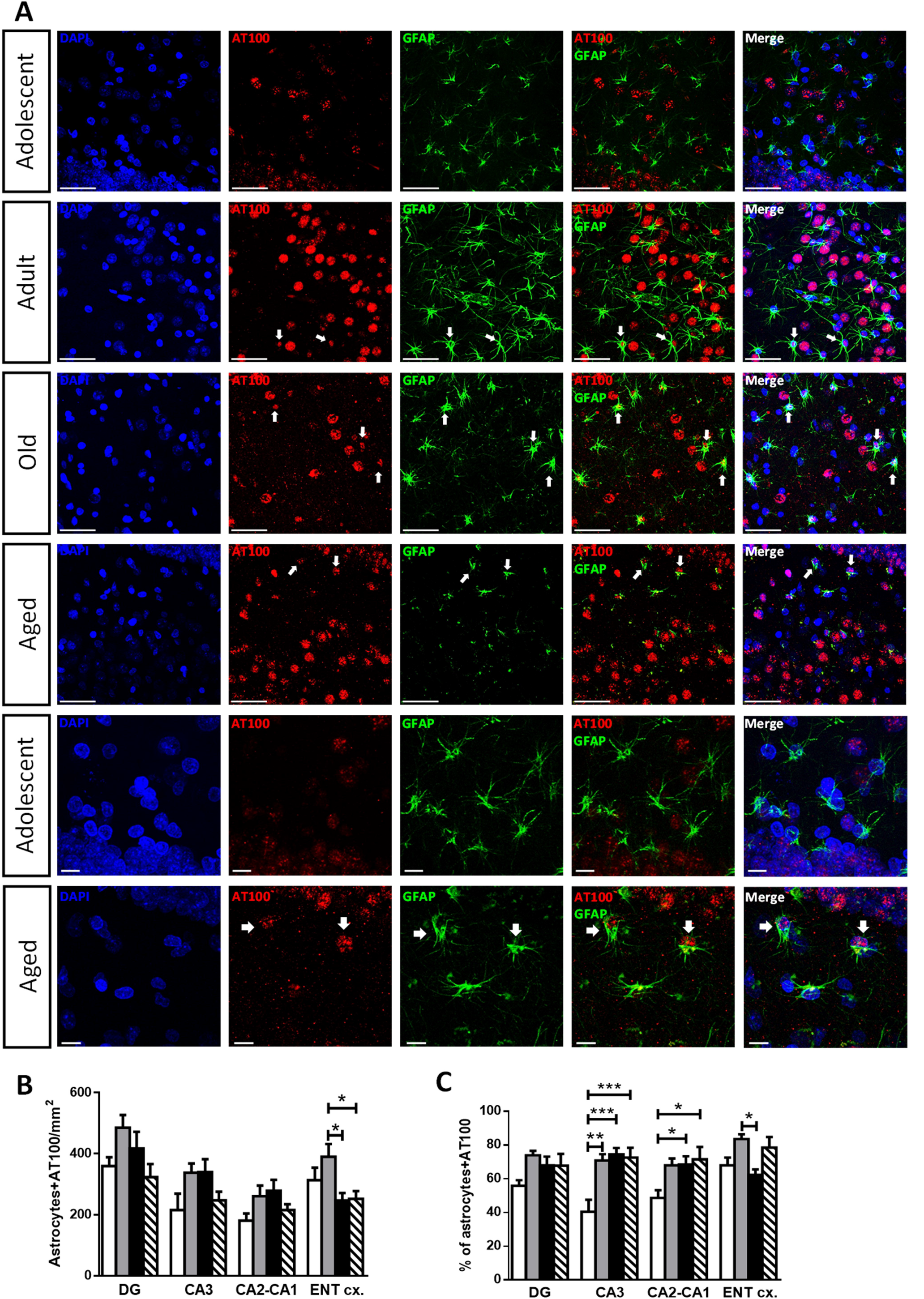
pTau in astrocytes of the marmoset. A) Representative photomicrographs of astrocytes (GFAP, in green) double labelled with AT100 (phosphorylation in the residues Thr212 and Ser214, in red) in CA3 of adolescent, adult, old and aged marmosets. In adolescents, few astrocytes were AT100+. In adult, old, and aged marmosets most astrocytes were AT100+. Note the intensity of AT100 labeling in the nucleus of astrocytes of aged subjects. DAPI was used as a nuclear counterstain. Scale bar 40 µm. **B)** Quantification of AT100+ astrocytes in different brain regions of the marmoset. DG, CA3 and CA2-CA1 regions did not differences in the amount of AT100+ astrocytes. ENT show a decreased number of AT100+astrocytes in old and aged marmosets, compared to adults. **C)** Percentage of AT100**+** astrocytes (number of AT100+ astrocytes / total number of astrocytes). There was an increase in the percentage of AT100+ astrocytes in CA3 region of adult, old and aged subjects compare to adolescents. In CA2-CA1 old and aged marmosets also showed a significant increase compared to adolescents. In ENT, the percentage of AT100+astrocytes was higher in adults compared to old subjects. Data represent means ± S.E.M. One-way ANOVA, Tukey post hoc analysis. *p < 0.05; **p < 0.01; ***p < 0.001.

### 3.6 S100A10+ astrocytes did not have an atrophic phenotype

In models of cerebral ischemia, astrocytes associated with a neuroprotective role are labeled with S100A10 protein (Zamanian et al., 2012). To determine if atrophic astrocytes lack S100A10 protein, we performed double immunolabeling: GFAP and S100A10. We decided to use aged marmosets, as this age group presents the largest number of atrophic astrocytes. Figure 6 shows representative images of S100A10- and S100A10+ astrocytes (panels A). It can be observed that in all regions analyzed, S100A10+astrocytes had longer processes than S100A10- astrocytes. The quantification of APL of S100A10- and S100A10+astrocytes demonstrate that Sholl curves of S100A10+ (yellow circles curves) and S100A10- (white squares curves) reach a maximum APL between radius 2 and 3 (figure 6B). In all regions analyzed, APL in S100A10+ astrocytes were higher than in S100A10- astrocytes, with significant differences in radius 2,3, 4, and 6 in DG; radius 4- 7 in CA3; radius 3,4 and 6 in CA2-CA1; radius 3 and 4 in ENT (Suppl. table 2). The quantification of total APL (figure 6C) demonstrates that S100A10+ astrocytes have significantly longer processes compared to S100A10- in CA3, CA2-CA1, and ENT. Concerning the quantification of the APV, S100A10+ astrocytes have significantly higher APV than S100A10- in DG, CA2-CA1, and ENT (figure 6D). Finally, the ABP was higher in S100A10+ in all regions, however, the differences were not significant (figure 6E).

**Figure 6.**
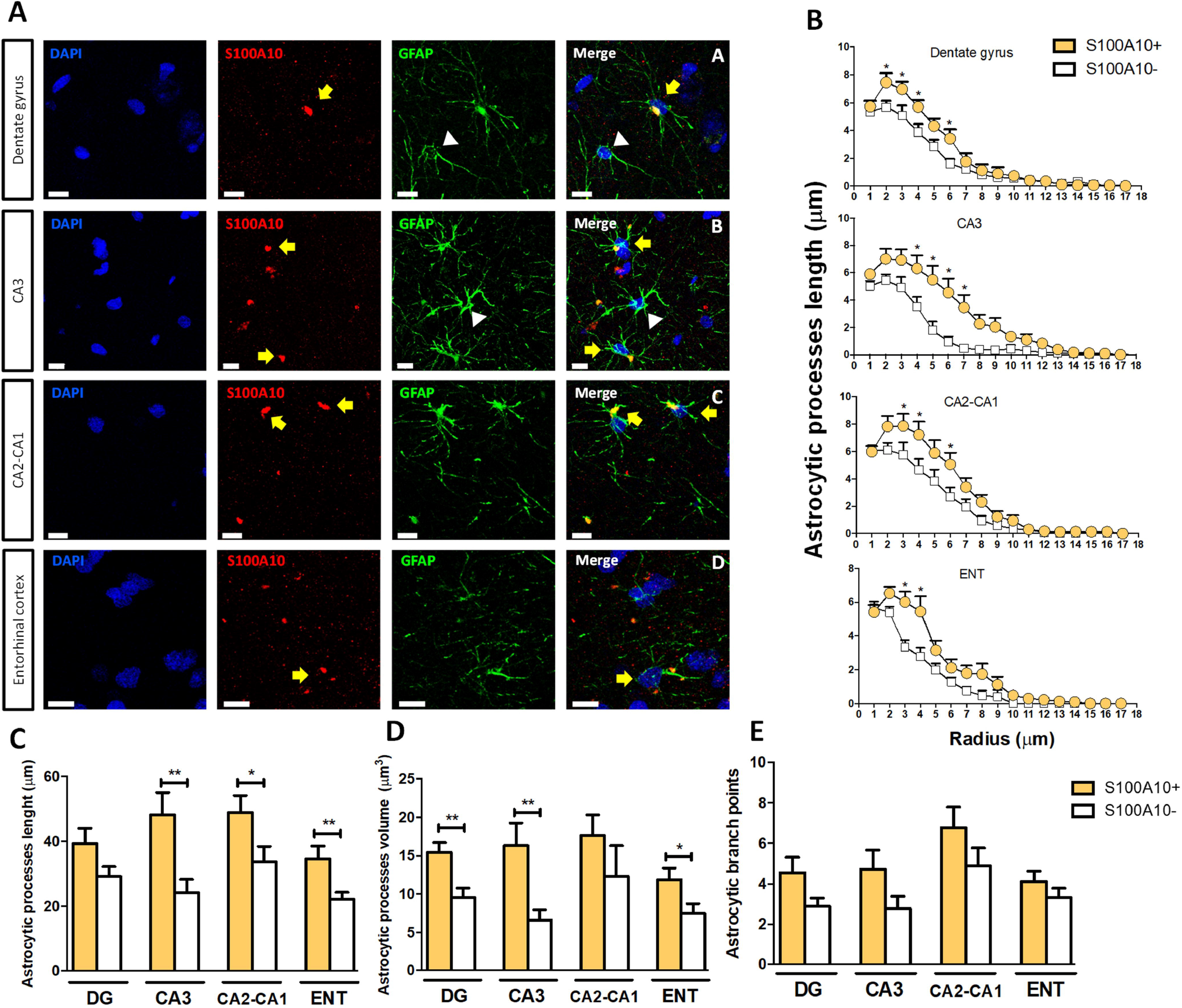
S100A10+ astrocytes had longer processes. A) Representative photomicrographs of astrocytes (GFAP+, green) with S100A10+ (red) in DG, CA3, CA2- CA1, and ENT of old marmosets. Most S100A10+ astrocytes (white arrowheads) had longer processes than S100A10- astrocytes (yellow arrows). DAPI was used as a nuclear counterstain. Scale bar 10 µm. **B)** Sholl analysis of S100A10+/- astrocytes. Astrocytic processes length was significantly higher in S100A10+ astrocytes than S100A10- astrocytes in DG (radius 2, 3, 4 and 6), CA3 (radius 4-7), CA2-CA1 (radius 3, 4 and 6), and ENT (radius 3 and 4). Multiple t-test followed by Holm-Sidak post hoc analysis. *p < 0.05. **C)** Quantification of the total APL of S100A10+ and S100A10-astrocytes. Total APL in CA3, CA2-CA1 and ENT was significantly higher in S100A10+ astrocytes than S100A10-. **D)** Quantification of APV. APV was significantly higher in S100A10+ astrocytes compared to S100A10- in DG, CA3 and ENT. **E)** Quantification of ABP. S100A10+ astrocytes presented larger ABP compared to S100A10-, but the differences were non-significant. Data represent means ± S.E.M. One-way ANOVA followed by Tukey post hoc analysis. *p < 0.05; **p < 0.01.

These results demonstrate that in aged marmoset brain, S100A10+ astrocytes are less prone to present atrophy, as longer astrocytic processes length and volume were observed in GFAP+/S100A10+ astrocytes, whereas astrocytes lacking S100A10 had distinctive morphological features of astroglial atrophy.

### 3.7 DNA fragmentation was present in S100A10- astrocytes of aged marmosets

To further determine if the atrophic phenotype and the lack of S100A10 were related to DNA damage, we used double-label immunofluorescence of GFAP and TUNEL reaction. We use this combination in the ENT, as this region showed a larger number of atrophic astrocytes. We first quantified the number of cells positive for TUNEL staining in adolescent, adult, old, and aged marmosets. There were cells positive for TUNEL in all age groups. However, the total number of cells with TUNEL increased in aged marmosets (figure 7A). Then, we determined if the number of GFAP+ and TUNEL+ cells changed during aging in the marmoset. We observed that adult marmosets had more TUNEL+ astrocytes compared to adolescents (figure 7B). In aged marmosets, more astrocytes were TUNEL+ with respect to the other age groups (figure 7C, white arrowheads). These results denote an increased number of cells with DNA fragmentation during aging. In particular, aged marmosets had the highest number of astrocytes with DNA fragmentation. This finding coincides with the abundant astrocytes with atrophic phenotype in aged marmosets.

**Figure 7.**
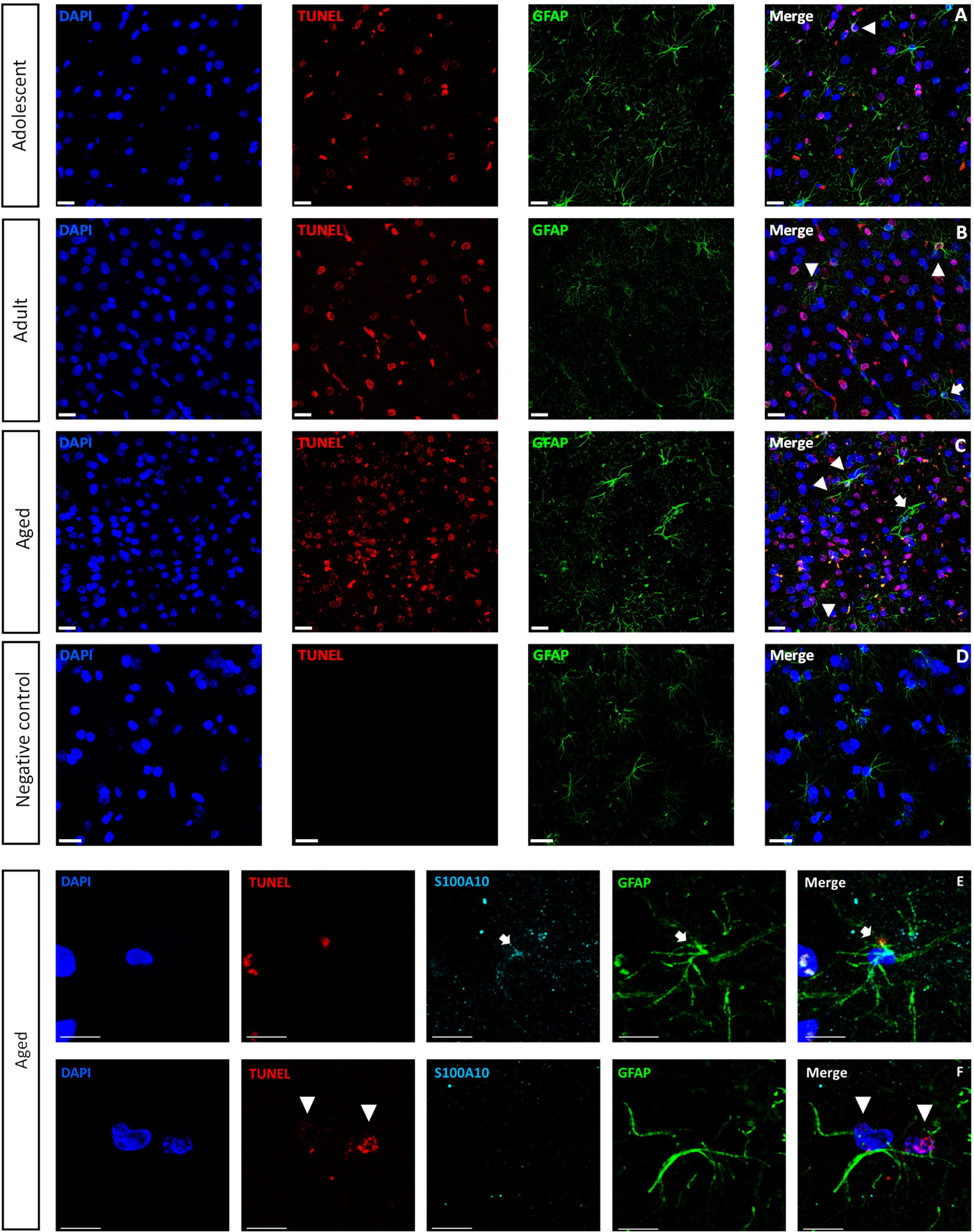
DNA fragmentation in S100A10- astrocytes. Representative photomicrographs of astrocytes (GFAP+, green) with DNA fragmentation (TUNEL, red) in the ENT of marmosets at different ages. In adolescents, most astrocytes do not present TUNEL, except for astrocytes with few small processes (white arrowhead). In adults, there were more TUNEL+ astrocytes (white arrowheads), however those astrocytes with longer processes were negative for TUNEL (white arrow). In aged marmoset, most astrocytes were TUNEL+ (white arrowheads). Note that TUNEL+ cells were higher in aged marmosets compared to adolescents and adults. Negative controls for TUNEL protocol are included. Lower two panels are representative images of the triple labeling: GFAP+ (green), S100A10+ (cyan) and TUNEL+ (red) in CA3 of aged marmosets. A magnification of S100A10+ astrocytes (white arrows) and two S100A10- astrocytes (white arrowheads) that were TUNEL+. Scale bar 20 µm for panels A-C and 10 µm for panels D and E. DAPI was used as a nuclear counterstain.

We aimed to determine if DNA fragmentation could be related to astrocyte atrophy (length of processes, branching points, and volume of dendrites). In addition, to understand if astrocyte atrophy could be related to a protective phenotype, we use S100A10 antibody in triple labeling (GFAP, S100A10, and TUNEL) in aged marmosets (figure 7D and E). Most S100A10+ astrocytes do not present TUNEL labeling in their nucleus. In contrast, S100A10- astrocytes showed clear TUNEL labeling. Moreover, TUNEL+ and S100A10- astrocytes present shorter processes and fewer branching points than S100A10+ astrocytes. These results suggest that the S100A10+ astrocytes are less vulnerable to DNA fragmentation and astrocyte atrophy during aging in marmosets.

## 4. Discussion

### 4.1 Increased number of reactive astrocytes in adult and old marmosets

In this study, we quantified the length, volume, and branch points of astrocytic processes from adolescents, adults, old, and aged marmosets in ENT and hippocampal region. We observed that in adult and old marmosets, astrocytes presented morphological features that resemble a reactive phenotype: increased length of processes and branching points compared to adolescent subjects (in DG and CA1-CA2). In contrast, aged marmosets presented shorter astrocytic processes in all regions analyzed, with morphological features of atrophic astrocytes. GFAP-FI increased in old marmosets in CA1-CA2 and CA3 regions compared to younger and older animals (Suppl. Figure 1). Despite the increased number of hypertrophic astrocytes in adult and old marmosets (increased processes length, volume, and branch points), the total number of astrocytes remains equal in all brain regions, except in DG (number of GFAP+ astrocytes were higher in adolescents vs. all ages) and ENT (number of GFAP+ astrocytes was lower in aged marmosets vs. all ages). Thus, our data indicate that adult and old marmosets develop mild astrogliosis, as astrocytes show hypertrophy and increased GFAP-FI, but without significant cellular proliferation.

Reactive astrogliosis is triggered in response to CNS injuries and diseases (Sofroniew and Vinters, 2010). It has been previously reported that old marmosets present amyloid plaques, pTau, enhanced concentrations of iron, RNA oxidation, and overactivation of microglia (Geula et al., 2002; Ridley et al., 2006; Rodriguez-Callejas et al., 2016; Rodríguez-Callejas et al., 2019; Sharma et al., 2019). All those factors may promote a reactive astrogliosis, as observed in the present study. It is important to note that CA2- CA1 region shows longer astrocytes in old marmosets compared to younger animals. These results coincide with our previous data, as CA2-CA1 is the region with the lowest number of 8OHG+ cells, in comparison with ENT; ENT had the lowest glia activation, but the highest number of 8OHG+ cells (Rodríguez-Callejas et al., 2019)(Rodríguez-Callejas et al., 2019). Therefore, hypertrophic astrocytes in old marmosets could reflect a protective mechanism.

### 4.2 Astroglial atrophy in the hippocampus of aged marmosets

In contrast to the hypertrophic phenotype observed in adult and old marmosets, aged subjects presented astrocytes with shorter process length, lower volume, and branch points, features of an atrophic phenotype, in all brain regions analized. Atrophy of GFAP+ astrocytes has been observed in ENT of aged mice (18 and 24 months)(Rodríguez et al., 2014). In 3xTg-AD mice, a mouse model of AD, GFAP+ astrocytes showed signs of atrophy in DG, CA1, ENT, and prefrontal cortex compared to control mice (Verkhratsky et al., 2015). The presence of atrophic astrocytes has been reported even in young 3xTgAD mice (1-month-old) in ENT compared to controls (Yeh et al., 2011). In brain samples of AD patients, a high number of atrophic astrocytes is observed, especially far from amyloid plaques (Matias et al., 2019). Contrary, hypertrophic astrocytes are found surrounding amyloid plaques (Matias et al., 2019). Pluripotent stem cells (iPSC)-derived astrocytes from Parkinson’s disease patients develop an atrophic phenotype with a decreased mitochondrial activity, ATP production, increase of glycolysis, and production of reactive oxygen species (ROS) (Ramos-Gonzalez et al., 2021).

The region with the most significant astrocytic alterations in the marmoset was ENT. In humans, ENT neurons are highly vulnerable to alterations in glucose concentrations and hypoxia (Fehm et al., 2006; Mattsson et al., 2016), and age-related mitochondrial dysfunction (Chen and Chan, 2009; Lin and Beal, 2006; Roselli and Caroni, 2015). Moreover, ENT neurons of layer II are the most vulnerable to age-related alterations (Stranahan and Mattson, 2010). Thus, our data may indicate that the abundance of atrophic astrocytes in ENT can be associated with the vulnerability of this brain region to the age-related metabolic and synaptic alterations. Therefore, the presence of atrophic astrocytes in hippocampus and ENT of aged marmosets indicates an ongoing neurodegenerative process beyond a normal aging process.

### 4.3 Abundant RNA oxidative damage in astrocytes of aged marmosets

In all regions analyzed, old and aged marmosets had a significantly increased number of astrocytes with RNA oxidation (8OHG+) compared to adolescents and adults. In a previous study, Bellaver and co-workers proved that hippocampal astrocytes of aged rats present high levels of ROS, oxidation of RNA, and a decreased mitochondrial membrane potential (Bellaver et al., 2017). Oxidative stress in astrocytes of aged rats can be induced by the overexpression of proinflammatory cytokines, increased expression and activity of proinflammatory enzymes, and decreased activity and expression of antioxidant enzymes (Bellaver et al., 2017). Oxidative damage alters the expression of multiple proteins important for the astrocytic function (Bellaver et al., 2017), culminating in the atrophy of these cells. Therefore, RNA oxidation might be associated with the atrophic phenotype in astrocytes of old and aged marmosets.

### 4.4 Nuclear pTau in astrocytes of adult, old and aged marmosets

In aging (Hof et al., 1996) and neurodegenerative diseases (i.e. AD, Down syndrome, and tauopathies) pTau causes its self-aggregation (Alonso et al., 1996, 2001, 2010; Despres et al., 2017; Liu et al., 2020) in straight and paired-helical filaments (PHF) which subsequently form the neurofibrillary tangles (NFT). In aged NHP, including the marmosets, pTau accumulation is observed in the hippocampus and cortex of old subjects (Darusman et al., 2014; Datta et al., 2021; Härtig et al., 2000; Oikawa et al., 2010; Paspalas et al., 2018; Perez et al., 2013; Rodriguez-Callejas et al., 2016; Schultz et al., 2000b, 2000a). In a previous study, we observed AT100 immunoreactivity mainly in the nucleus of principal neurons from the granular layer and polymorphic layers of DG, and *str. pyramidale* of CA3 and CA2-CA1 in adult, old, and aged marmosets (Rodriguez-Callejas et al., 2016). In this study, we detected nuclear pTau (AT100+) in GFAP+ astrocytes in ENT, CA3 and CA2-CA1. The percentage of GFAP+ astrocytes with nuclear pTau increased in adults, old and aged marmosets in DG, CA2-CA1, compared to adolescents.

*In vitro* studies demonstrate that tau binds nuclear DNA (single and double-stranded) (Hua et al., 2003; Krylova et al., 2005; Padmaraju et al., 2010) by the proline-riched domain and the microtubule-binding domain (Qi et al., 2015; Wei et al., 2008). DNA-tau binding increases the stability of the chromosomes (Camero et al., 2014; Rossi et al., 2008; Sjöberg et al., 2006) and protects the DNA against oxidative stress and hyperthermic conditions (Sultan et al., 2011; Wei et al., 2008). Diverse studies report the presence of nuclear pTau using AT100 antibody in fibroblast cultures (Rossi et al., 2008), rats (Gärtner et al., 1998), tree shrews (Rodriguez-Callejas et al., 2020), and humans (Gil et al., 2017; Hernández-Ortega et al., 2016). In human studies, the number of AT100+ cells and the intensity of the label increase in the granular layer of DG and the *str. pyramidale* of CA1 in cognitive-healthy old subjects compared to adolescents and adults (Gil et al., 2017). However, in AD patients, the number of AT100+ cells significantly decreases as the disease progresses, indicating that the protective role of this pTau has been exhausted (Gil et al., 2017). In the current study, we observed an increased number of astrocytes with nuclear pTau (AT100) with aging in the hippocampus. However, in ENT, the number of astrocytes with nuclear pTau was decreased in aged marmosets. The reduction in the number of astrocytes with nuclear pTau can indicate a loss of its nuclear protective mechanism, similar to the late stages of AD (Gil et al., 2017).

### 4.5 Lack of S100A10 protein associates with atrophic astrocytes

Lipopolysaccharide exposure and inflammatory insults cause the release of proinflammatory cytokines by reactive astrocytes (proinflammatory-phenotype) (Zamanian et al., 2012). Contrarily, reactive astrocytes can release neurotrophic factors in conditions such as ischemia, promoting tissue repair and recovery; these astrocytes are termed “neuroprotective-astrocytes” (Zamanian et al., 2012). Neuroprotective astrocytes can actively clear Aβ and degrade it, protecting neurons against their neurotoxic effects (Xiao et al., 2014). In normal aging, proinflammatory astrocytes are the predominant type in the hippocampus and striatum of rodents (Clarke et al., 2018). In advanced stages of AD, reactive astrocytes lose their neuroprotective role, and instead, they release proinflammatory cytokines provoking neuronal damage and synapse degeneration (Lue et al., 1996; Perez-Nievas et al., 2013).

In old mice, astrocytes in the hippocampus and striatum show an up-regulation of genes related to the proinflammatory phenotype rather than neuroprotective-related genes. Moreover, in old animals, the number of astrocytes that express a proinflammatory marker (*C3*) is larger than astrocytes that express a neuroprotective specific gene (*Emp1*) (Clarke et al., 2018). In diverse neurodegenerative diseases, such as AD, Parkinsońs disease, Huntington, amyotrophic lateral sclerosis, and multiple sclerosis, astrocytes have a proinflammatory profile (Liddelow et al., 2017). The increased number of proinflammatory astrocytes during aging and neurodegeneration may cause inflammation by releasing inflammatory cytokines and complement components (Jang et al., 2013; Liddelow et al., 2017). Contrary, neuroprotective-type astrocytes express S100A10 protein (Clarke et al., 2018). Here, we observed that S100A10+ astrocytes had larger processes than S100A- astrocytes, in all regions analyzed. This may indicate that S100A10 protein confers neuroprotection against atrophy. We did not use a marker of proinflammatory astrocytes, that could help to determine whether this shift in morphology and lack of S100A10 protein is associated with a specific reactive state. Thus, future studies are needed to determine better the molecular profile of the reactive astrocytes in the selected brain regions of the marmoset during the aging process.

### 4.6 DNA fragmentation is found in S100A10- astrocytes

Apoptosis is a mechanism of regulated cell death that occurs under normal physiological conditions but also plays a crucial role in diverse pathologies (Bertheloot et al., 2021; Xu et al., 2019). Apoptosis induces morphological and biochemical changes in the cells, such as cell and nuclear shrinkage, chromatin condensation (pyknosis), DNA fragmentation, and membrane-bound cell fragments (apoptotic bodies) (Majtnerová and Roušar, 2018; Xu et al., 2019). DNA fragmentation is the main feature of apoptosis and can be determined by TUNEL staining (Arends et al., 1990; Walker et al., 1994). In marmosets, we observed an increased number of TUNEL+ nuclei in aged subjects compared to adolescents and adults (Figure 7). In aged marmosets, most astrocytes were TUNEL+, coinciding with the significant increase of atrophic astrocytes. To further understand this correlation, we labeled GFAP+ astrocytes with S100A10 and TUNEL in the hippocampus of aged marmosets. Most S100A10- astrocytes were positive for TUNEL staining. On the other hand, most of the S100A10+ astrocytes did not show TUNEL labeling. These results suggest that S100A10+ protein protects astrocytes from presenting DNA fragmentation and an atrophic phenotype.

## Conclusion

Our results show that adult and old marmosets had a reactive astrogliosis in the hippocampus and ENT compared to adolescent and aged marmosets. However, aged marmosets show morphological alterations in all regions analyzed, as they present a prominent atrophic phenotype. In addition, damage to RNA was observed in astrocytes of old and aged marmosets compared to younger animals. Nuclear pTau was detected in astrocytes of all regions analyzed showing an age-dependent increase in hippocampal regions. However, in ENT the number of astrocytes with nuclear pTau was reduced in aged marmosets. Furthermore, neuroprotective-type astrocytes (S100A10+) had an elongated morphology (length, volume, and branch points; hypertrophic) than S100A10+ astrocytes (atrophic). Neuroprotective-type astrocytes (S100A10+) did not presented DNA fragmentation, whereas lack of S100A10 associated with DNA damage. Here we show alterations in astrocytes’ activation during aging in marmosets, with an enhanced reactivity in adult and old animals, but morphological modifications (atrophy) in aged animals that go beyond normal aging. Thus, our data contribute to the growing body of literature underpinning the use of the marmoset as a suitable animal model to investigate the etiological factors and processes associated with neurodegeneration.

## ACKNOWLEDGMENTS

Rodriguez-Callejas, J.D. CONACYT Scholarship no. 308515

## SUPPLEMENTARY INFORMATION

**Supplementary Figure 1.**
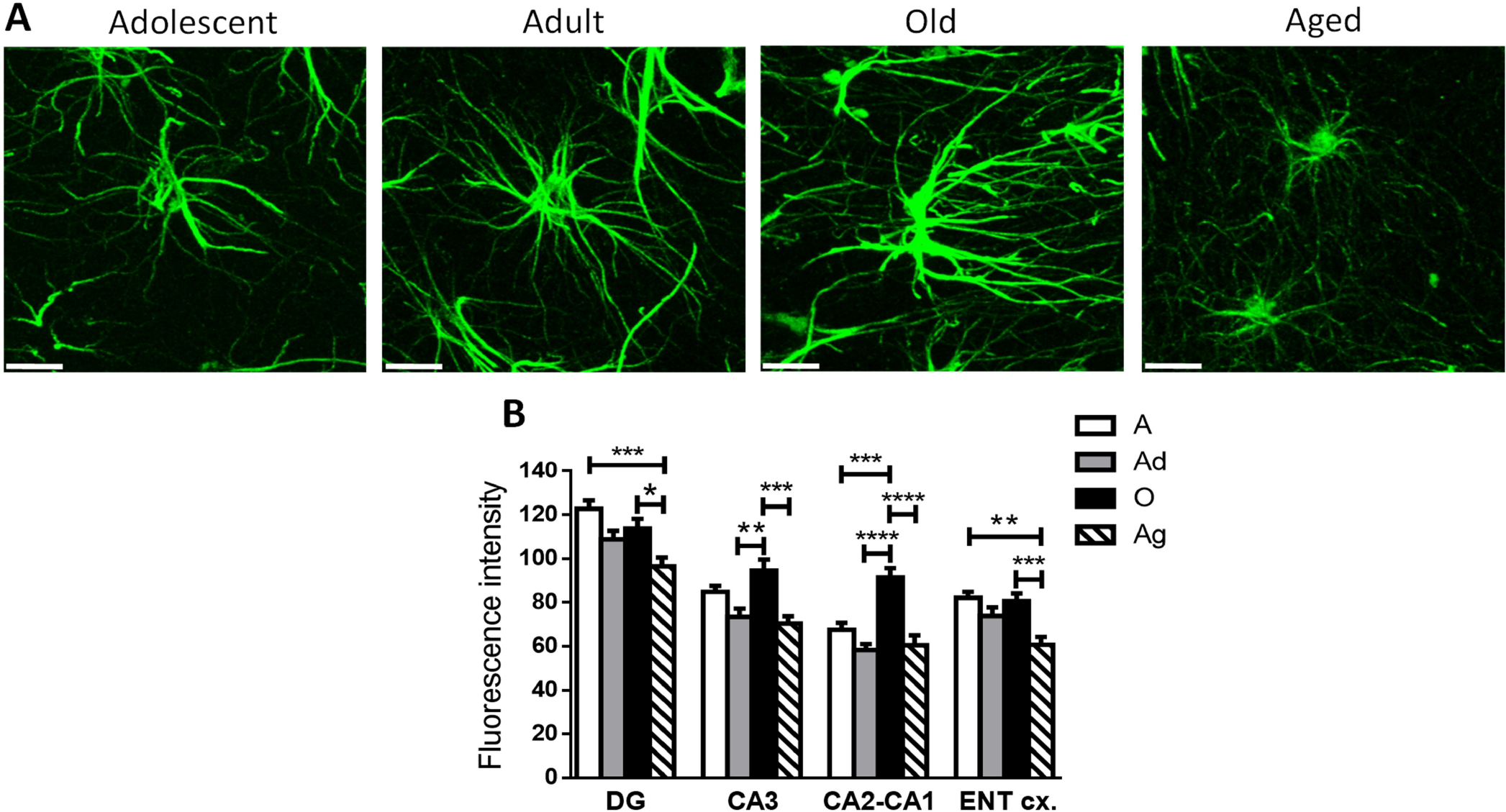
GFAP fluorescence intensity (FI) of astrocytes in different brain regions during aging. A) Representative photomicrographs of GFAP+ astrocytes of adolescent, adult, old and aged marmosets. Scale bar 10 µm. **B)** Quantification of fluorescence intensity of GFAP+ astrocytes located in the hippocampus (DG, CA3 and CA2-CA1 regions) and the ENT. In DG and ENT, aged marmosets showed a decreased FI compared to adolescent and old marmosets. In CA3 and CA2-CA1, old subjects showed an increased FI than adolescent, adult and aged marmosets. Data represent means ± S.E.M. One-way ANOVA, Tukey post hoc analysis. *p < 0.05; **p < 0.01; ***p < 0.001; ****p < 0.0001.

